# Variation in competitive ability with mating system, ploidy and range expansion in four *Capsella* species

**DOI:** 10.1101/214866

**Authors:** Xuyue Yang, Martin Lascoux, Sylvain Glémin

## Abstract

This preprint has been reviewed and recommended by Peer Community In Evolutionary Biology (https://dx.doi.org/10.24072/pci.evolbiol.100054)

Self-fertilization is often associated with ecological traits corresponding to the ruderal strategy in Grime’s Competitive-Stress-tolerant-Ruderal (CSR) classification of ecological strategies. Consequently, selfers are expected to be less competitive than outcrossers, either because of a colonization/competition trade-off or because of the deleterious genetic effects of selfing. Range expansion could reduce further competitive ability while polyploidy could mitigate the effects of selfing. Although suggested by meta-analyses, these predictions have not been directly tested yet. We compared the competitive ability of four *Capsella* species differing by their mating system and ploidy level. For vegetative traits we found no difference in competitive ability neither among species nor among populations. For flower production, we found that the two diploid selfing species (*C. rubella* and *C. orientalis*) were more sensitive to competition than the diploid outcrosser (*C. grandiflora*), and that the tetraploid selfer (*C. bursa-pastoris*) was intermediate. Within *C. bursa-pastoris*, we also found that sensitivity to competition increased in parallel to range expansion. These results highlight the possible roles of ecological context and ploidy in the evolutionary trajectories of selfing species.

## Introduction

The transition from outcrossing to selfing is very common in flowering plants. It is likely that many shifts to selfing abort early because of the rapid deleterious effect of selfing (Willi 2013; Griffin & Willi 2014; Abu Awad & Billiard 2017). But, if successful, the evolution of selfing is often associated with habitat shift and range expansion, thanks to the reproductive assurance offered by the ability to self (Randle *et al.* 2009; Grossenbacher *et al.* 2015). Selfers are more frequently found under conditions in which obligatory outcrossers pay the demographic cost of mate limitation, such as disturbed and patchy habitats or newly available environments (Baker 1967; Munoz *et al.* 2016). In addition to reproductive traits, often referred to as the “selfing syndrome” (Sicard & Lenhard 2011), other traits may thus evolve after the transition to selfing in relation with new ecological conditions. For instance, selfing is often associated with weedy habit (Clements *et al.* 2004) and invasiveness (van Kleunen *et al.* 2008). More generally, when referring to Grime’s Competitive-Stress-tolerant-Ruderal (CSR) ecological strategies (Grime 1974), an excess of selfers is observed among ruderal species, corresponding to early colonizers in ecological successions, whereas more competitive species tend to be outcrossers (Munoz *et al.* 2016).

At least two mechanisms could explain the negative association between selfing and competitiveness. First, a colonization/competition trade-off could be involved, where selection for better colonizing ability would be at the cost of lower competitive ability (Burton *et al.* 2010). Related to this hypothesis, if costly, traits involved in competitive ability could be selected against during range expansion when low competitive environments are colonized (Bossdorf *et al.* 2004). Alternatively, selfing could affect competitive ability through its negative genetic effects, especially the accumulation of weakly deleterious mutations. Deleterious mutations affecting the efficiency of resource acquisition are predicted to reduce competitive ability (Agrawal 2010; Agrawal & Whitlock 2012). Selfing species are expected to rapidly purge inbreeding depression caused by strongly deleterious recessive alleles but can accumulate weakly deleterious mutations on the long term because of linked selection effects (Wright *et al.* 2008; Glémin & Galtier 2012). From a demographic perspective, the purging of inbreeding depression can help survive bottlenecks associated with initial colonisation of new habitats but weakly deleterious mutations can also accumulate later on during demographic expansion, the so-called “expansion load” (Peischl *et al.* 2013; Peischl & Excoffier 2015). In agreement with a possible effect of deleterious mutations on competitive ability, inbred individuals have been found to suffer more from competition than outbred ones (Cheptou *et al.* 2000; Cheptou *et al.* 2001; Yun & Agrawal 2014). Polyploidy is another factor that can alter the above predictions. Polyploidy is often associated with selfing/self-compatibility because polyploidy can result in a breakdown of self-incompatibility systems and/or because polyploids can establish more easily if they can self, avoiding the detrimental effect of crossing with their diploid progenitors (Barringer 2007; Robertson *et al.* 2011). Polyploidy is supposed to increase competitiveness (Comai 2005; te Beest *et al.* 2012) and to reduce the genetic load, at least for a transient period after the formation of the polyploid species (Otto & Whitton 2000). Thus, polyploidy could buffer the deleterious effects of selfing. According to the rationale presented above, and whatever the underlying causes, we predict that (i) transition from outcrossing to selfing should lead to reduced competitive ability, (ii) this effect should be less pronounced in polyploid than in diploid selfers, and (iii) within species, competitive ability should decline during range expansion.

The *Capsella* genus (Brassicaceae) is a good model to address these questions. It comprises four closely related annual species with contrasting mating systems and ploidy levels but other life-history and morphological traits are very similar, including growth habit, phenology, seed size and shape, and dispersal mode. They also live in similar open and disturbed habitats. The only outcrossing species in the genus, *C. grandiflora* (Fauché & Chaub.) Boiss. is restricted to western Greece and Albania but can be found in very large populations in agricultural and disturbed environments. It has a large effective population size, with strong evidence of efficient positive and purifying selection (Foxe *et al.* 2009; Slotte *et al.* 2010; Williamson *et al.* 2014). The three selfing species include two diploids: *C. orientalis* Klokov diverged from *C. grandiflora* about one million years ago and *C. rubella* Reut. much more recently (30,000–50,000 years ago Foxe et al., 2009, Guo et al., 2009). Selfing thus evolved twice independently. Both species have much larger species range than *C. grandiflora* but also much patchier distributions. Their effective population sizes are much smaller and both species have accumulated weakly deleterious mutations (Foxe *et al.* 2009; Guo *et al.* 2009; Slotte *et al.* 2013; Douglas *et al.* 2015; Kryvokhyzha *et al.* submitted). The third selfing species, *C. bursa-pastoris* (L.) Medik., is an allo-tetraploid with disomic inheritance that originated about 200,000 years ago as a hybrid between *C. orientalis* and *C. grandiflora* (Douglas *et al.* 2015). It is not known whether selfing was inherited from the *C. orientalis* parent or whether it evolved a third time independently. *C. bursa-pastoris* has by far the largest range with an almost worldwide distribution owing to historical colonization in Eurasia and very recent human dispersal across other continents (Cornille *et al.* 2016). It also has a patchy distribution, especially in urban areas. In Greece, *C. grandiflora*, *C. rubella* and *C. bursa-pastoris* can be found in sympatry (M.L. and S.G. personal observation). In Eurasia, three *C. bursa-pastoris* genetic clusters can be distinguished, likely corresponding to colonization events from Middle East (the probable center of origin) to Europe and then to Eastern Asia. Across its expansion range, genetic diversity decreases and deleterious mutations accumulates from Middle East to Eastern Asia (Cornille *et al.* 2016; Kryvokhyzha *et al.* submitted), in agreement with population genetic predictions on range expansion dynamics (Excoffier *et al.* 2009; Peischl *et al.* 2013; Peischl & Excoffier 2015). During expansion, and especially in the expansion front in China, *C. bursa-pastoris* showed signature of introgression by its selfing parent, *C. orientalis*,

In a previous study (Petrone Mendoza *et al.* 2018), *C. rubella* has been shown to be more sensitive to competition than both *C. grandiflora* and *C. bursa-pastoris*, but no significant difference was observed between *C. grandiflora* and *C. bursa-pastoris*. However, this initial study focused only on Greek populations where the three species are found in sympatry. It was not representative of the whole species range of *C. bursa-pastoris* and one of the diploid progenitors of *C. bursa-pastoris*, *C. orientalis,* was missing. Here, we performed a new competition experiment to compare the competitive ability of *C. bursa-pastoris* with its two parental species, *C. grandiflora* and *C. orientalis*, and within *C. bursa-pastoris*, among populations across the species range. For completeness and comparison with Petrone Mendoza et al. (Petrone Mendoza *et al.* 2018) we also added *C. rubella*. Our working hypotheses were (i) that competitive ability of *C. bursa-pastoris* should be intermediate between *C. grandiflora* on the one hand, and *C. rubella* and *C. orientalis*, on the other hand, and (ii) that competitive ability within *C. bursa-pastoris* should decline in parallel to range expansion.

## Material and methods

### Studied species and sampling

We sampled the four *Capsella* species: the diploid outcrosser, *C. grandiflora*, the two diploid selfers, *C. rubella* and *C. orientalis* and the tetraploid selfer, *C. bursa-pastoris*. One accession was initially sampled from each of the 62 populations of *C. bursa-pastoris* studied by Cornille et al. (Cornille *et al.* 2016), including 22 European, nine Middle Eastern, and 31 Chinese populations, respectively. We added five populations from Central Asia, an area that was not sampled by Cornille et al. (Cornille *et al.* 2016) (Table S1). In addition, 15 to 35 accessions from each of the three other *Capsella* species sampled over their species range were used for interspecific comparisons, corresponding to 5 *C. orientalis*, 16 *C. rubella* and 9 *C. grandiflora* populations (Table S1).

A previous work (Petrone Mendoza *et al.* 2018) has shown that the difference in sensitivity to competition was weakly affected by the nature of the competitor. As a result we used only one, non-*Capsella*, competitor species to simplify the experimental design and ensure the same inter-specific competition for the four species. As in Petrone Mendoza et al. (Petrone Mendoza *et al.* 2018), we used *Matricaria chamomilla*, an annual Asteraceae species, which was found to co-occur with *Capsella* species in Greek populations and at least with *C. bursa-pastoris* in other European populations (personal observations). More generally, *Capsella* species and *M. chamomilla* have overlapping distributions (Europe and temperate Asia), and share their rosette growth pattern and weedy habits. We used commercial seeds for *M. chamomilla*, ensuring good germination and homogeneity among plants.

### Experimental design

A competition experiment was performed in order to assess the competitive ability of all four *Capsella* species. Each accession was tested with four replicates with and without competition in a complete random block design. For the “competition” treatment, one focal *Capsella* individual was sown in the middle of an 11cm x 11cm x 11cm pot, surrounded by four *M. chamomilla* competitors. For the “alone” treatment, focal individuals were sown without competitors.

30 seeds from each accession were first surface-sterilized. For stratification, seeds were sown in agar plates in darkness for 6 days at 4°C. Following sterilization and stratification, agar plates were moved to a growth chamber at 22 °C with 16h light and 8h darkness, under a light intensity of 130 lmol/m2/s for germination. After 5 days, germinated competitors were transferred to pots filled with standard culture soil and were placed randomly in two growth chambers (each growth chamber has two blocks) under the same temperature and light conditions as during the germination period. After recording the germination rates of each accession, focal species were sown five days after the competitors in order to ensure strong enough competition against *Capsella* species. Accessions with less than eight germinated seeds (the minimum required to fulfil the design) were removed. Seedlings were watered daily for 2 weeks and every other day until the end of the experiment.

A series of fitness-related traits of focal individuals were measured during the experiment. Three weeks after the seedlings had been sown, two perpendicular diameters of the rosette were measured a first time and then again one week later. We used the product of these two diameters as a proxy for rosette surfaces at time t1 and t2, respectively (S1 and S2). From them, we calculated growth rate as the relative difference of rosette surface: (S2 – S1)/S1. When plants senesced, approximately 40 days after the beginning of flowering, the total number of flowers was counted for each focal individual. As in Petrone Mendoza et al. (Petrone Mendoza *et al.* 2018), we used flower numbers, rather than fruit or seed numbers, to allow comparison between the four species because *C. grandiflora*, which is self-incompatible, set almost no fruit due to the absence of pollinators in the growth chambers. All measurements were done by a single person (X. Y.).

### Genetic diversity data

Except for the five recently collected individuals from Central Asia, population structure of *C. bursa-pastoris* has been inferred using Genotyping By Sequencing (GBS) data in Cornille et al. (Cornille *et al.* 2016). In our analysis, we used the same SNP data to estimate genetic diversity. In brief, 253 accessions from the 62 populations throughout Europe, Middle East and Asia were genotyped by sequencing (GBS) using the *C. rubella* genome v1.0 (Slotte *et al.* 2013) as a reference for mapping. After checking for sequencing reliability, filtering against sequencing errors and fixed heterozygote sites, 4274 SNPs were available for further analysis (see Cornille *et al.* 2016 for details). Mean pairwise nucleotide differences (π) (Nei 1978) were obtained with *arlsumstat* (Excoffier & Lischer 2010) for each sampling site with 0.5% missing data allowed. π values were corrected for missing data and divided by 64-bp (GBS markers’ length) to obtain the mean pairwise nucleotide differences per site, which we used as a proxy for local genetic drift. Genetic diversity data are given in Table S2.

### Data analysis

Data were analysed with generalized linear mixed models in R version 3.3.2 (R Development Core Team 2011). For all variables, block, species, treatment and all pairwise interactions were included as fixed effects and accessions as random effect. For reproductive traits we also added rosette surface at time t2 and its interactions with the other variables as fixed effects. For analyses within *C. bursa-pastoris* we used the area of origin as a factor instead of species. Interactions were tested first; significant interactions were then kept to test for main effects (type III ANOVA). For linear models, we used the *anova* function of the standard R distribution. For generalized linear models we performed analyses of deviance using the *Anova* function of the *car* package using the type III ANOVA option (Fox & Weisberg 2011). For linear models, random effects where tested by likelihood ratio tests using the *rand* function of the *lmerTest* package (Kuznetsova *et al.* 2016). For generalized linear models we also ran their equivalent models without random effects and compared the likelihood of the models with the *anova* function.

Rosette surface and growth rate were analysed using a mixed linear model fitted by maximum likelihood with the *lmer* function of the *lme4* package (Bates *et al.* 2015). The flowers number distribution was bimodal with a mode at 0 and another around 1000 (see results). Thus it was analysed in two steps. First, we analysed the proportion of flowering plants with a binomial model and a logit link using the *glmer* function of the R package *lme4* (Bates *et al.* 2015). Dead plants were included in the non-flowering category. Second, we excluded plants that did not flower and analysed flower number with a negative binomial model and a log link with the *glmmadmb* function of the *glmmADMB* package (Fournier *et al.* 2012; Skaug *et al.* 2013).

For the comparison among species, we were specifically interested in the species x treatment interaction term to determine whether the effect of competition differed among species. For comparison within *C. bursa-pastoris*, we were interested in the area x treatment interaction term to determine whether the effect of competition differed across the expansion range. If these interaction terms were significant, differences among interaction components (“contrast of contrasts”) were tested with the *testInteraction* function of the *phia* package (https://CRAN.R-project.org/package=phia), with False Discovery Rate (FDR) correction for multiple testing. To avoid over-parameterization, other interaction terms were removed if not significant before contrast testing. To get a more direct estimate and more intuitive interpretation of the result, we computed a competition index, as defined in Petrone Mendoza et al. (Petrone Mendoza *et al.* 2018):

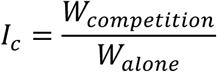

where *W_alone_* and *W_competition_* are the least-square mean of a trait without and with competition, respectively. Mean effects were estimated with the *lsmean* function of the *lmerTest* package (Kuznetsova *et al.* 2016). The lower *Ic*, the more sensitive to competition the species or the geographic group is.

Finally, in *C. bursa-pastoris,* we tested whether the area x treatment interaction could be explained by differences in genetic diversity among populations, with the prediction that the effect of competition should be more severe in low polymorphic populations. To do so, we ran a simple linear model with block and mean pairwise nucleotide differences (π), as fixed effects, π replacing the area effect. As each population is represented by its genetic diversity we did not include populations as random effect in this model. The five accessions from central Asia were not included in this analysis because we lack genetic data for them. The dataset and the R script used for analysis and plotting are provided as supplementary files.

## Results

After having discarded accessions with too few seeds, the final data set contained 13 accessions of *C. grandiflora*, 33 of *C. rubella*, 19 of *C. orientalis* and 49 of *C. bursa-pastoris*, distributed in the four geographic regions as follows: nine from the Middle East, 17 from Europe – including eastern Russian populations that belong to the same genetic cluster, see (Cornille *et al.* 2016) – five from central Asia – without genetic characterization – and 18 from China (Table 1). During the experiment survival rates were very high (> 90%) for all species and most plants flowered (Table 1).

**Table 1:** characteristics of the dataset.

| Species |  | # accessions used | Survival rate (%) | Flowering rate (%) |
| --- | --- | --- | --- | --- |
| *C. grandiflora* |  | 13 | 96.8 | 87.5 |
| *C. rubella* |  | 33 | 92.1 | 78.4 |
| *C. orientalis* |  | 19 | 100 | 100 |
| *C. bursa-pastoris* | Total | 49 | 91.1 | 90.1 |
|  | Middle East | 9 | 100 | 100 |
|  | Europe | 17 | 95.6 | 95.6 |
|  | Central Asia | 5 | 90.6 | 87.5 |
|  | China | 18 | 82.6 | 82.6 |

We first compared the four species, considering *C. bursa-pastoris* as a single unit. Rosette surfaces at the two times, but not growth rate, were significantly and negatively affected by competition (Table 2) but the effect was modest and competition only reduced rosette surface by around 5% (Figure 1). Moreover, the four species did not differ in their sensitivity to competition (the species x treatment effects were not significant, Table 2 and Figure 1). On the contrary we detected a strong effect of competition on flowers number and the four species differed in their sensitivity to competition (Table 3). The differences among species can be summarized by the competition index (Figure 1) and tested by pairwise contrasts on the species x treatment interactions (Table S3). *C. orientalis* and *C. rubella* were the most affected by competition with a reduction in flowers number by a factor two: *Ic* = 0.52 for both species whereas *C. grandiflora* was the least affected: *Ic* = 0.74. *C. bursa-pastoris* was intermediate with *Ic* = 0.65. The contrast analysis on interactions showed that all pairwise species comparisons are significant except between *C. orientalis* and *C. rubella* (Table S3).

**Figure 1:**
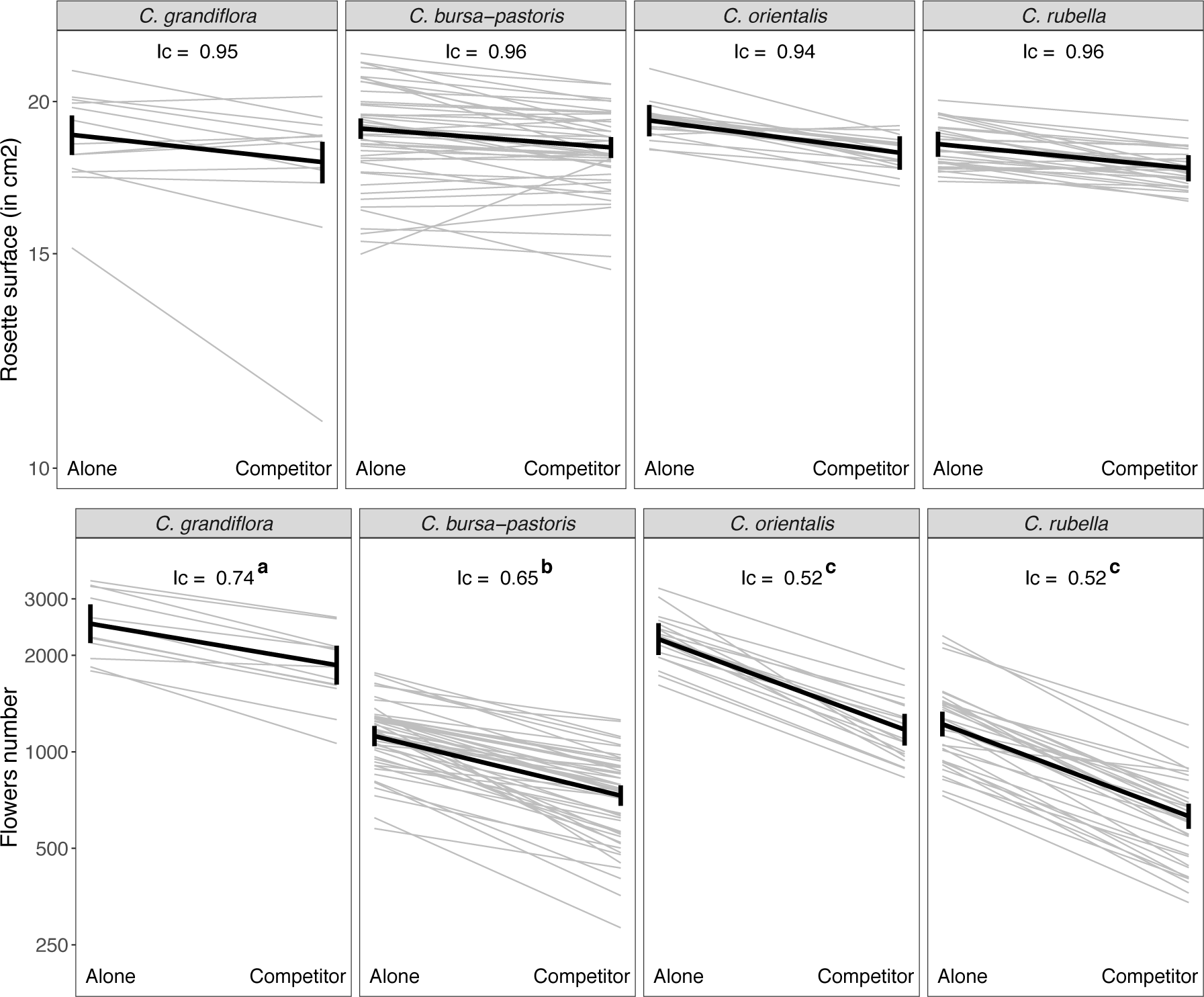
Rosette surface and flowers number without and with competitors in the four species. Each grey line corresponds to one accession (average over the four blocks) and the black lines join the least;square mean estimates (with confidence intervals). Ic: competition index. Ic with different letters corresponds to significant treatment x species interactions (see Tables S3 to S5).

**Table 2.**
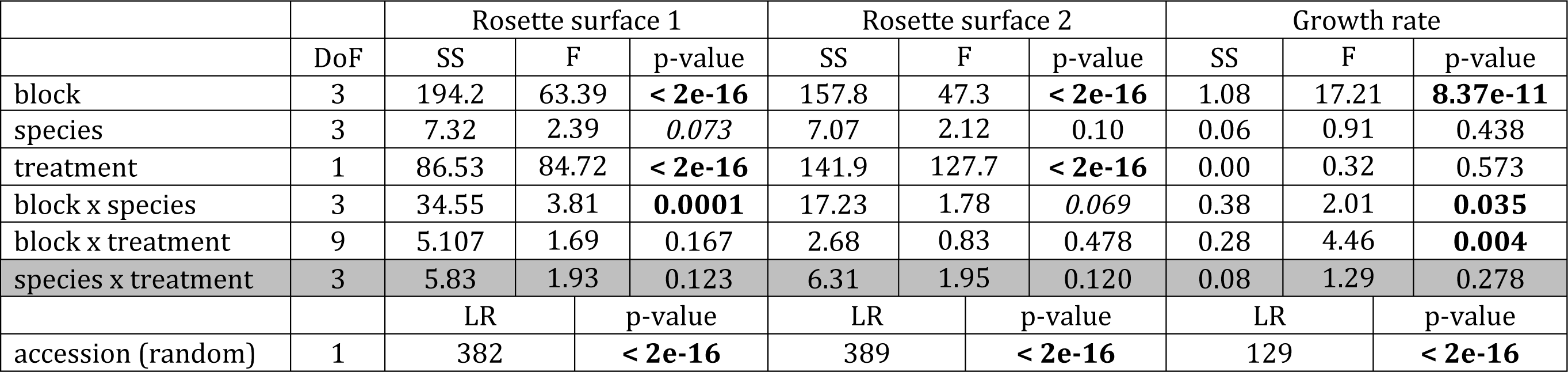
Analyses of variance for vegetative traits for the comparison among species. p-values ≤ 0.05 are in bold and p-values ≤ 0.1 in italics. The grey line corresponds to the interaction term of interest. DoF, and LR stands for Degrees of Freedom and Likelihood Ratio, respectively.

|  |  | Rosette surface 1 |  |  | Rosette surface 2 |  |  | Growth rate |  |  |
| --- | --- | --- | --- | --- | --- | --- | --- | --- | --- | --- |
|  | DoF | SS | F | p-value | SS | F | p-value | SS | F | p-value |
| block | 3 | 194.2 | 63.39 | **< 2e-16** | 157.8 | 47.3 | **< 2e-16** | 1.08 | 17.21 | **8.37e-11** |
| species | 3 | 7.32 | 2.39 | *0.073* | 7.07 | 2.12 | 0.10 | 0.06 | 0.91 | 0.438 |
| treatment | 1 | 86.53 | 84.72 | **< 2e-16** | 141.9 | 127.7 | **< 2e-16** | 0.00 | 0.32 | 0.573 |
| block x species | 3 | 34.55 | 3.81 | **0.0001** | 17.23 | 1.78 | *0.069* | 0.38 | 2.01 | **0.035** |
| block x treatment | 9 | 5.107 | 1.69 | 0.167 | 2.68 | 0.83 | 0.478 | 0.28 | 4.46 | **0.004** |
| species x treatment | 3 | 5.83 | 1.93 | 0.123 | 6.31 | 1.95 | 0.120 | 0.08 | 1.29 | 0.278 |
|  |  | LR |  | p-value | LR |  | p-value | LR |  | p-value |
| accession (random) | 1 | 382 |  | **< 2e-16** | 389 |  | **< 2e-16** | 129 |  | **< 2e-16** |

**Table 3.** Analyses of deviance for reproductive traits for the comparison among species. Same legend as Table 2.

|  | DoF | Proportion of flowering plants |  | Number of flowers |  |
| --- | --- | --- | --- | --- | --- |
|  |  | LR | p-value | LR | p-value |
| block | 3 | 3.59 | 0.309 | 38.61 | **2.10e-08** |
| rosette surface at t<sub>2</sub> | 1 | 2.89 | 0.089 | 1.12 | 0.290 |
| species | 3 | 18.70 | **0.0003** | 180.9 | **< 2.2e-16** |
| treatment | 1 | 3.41 | 0.065 | 23.19 | **1.47e-06** |
| block x rosette surface at t<sub>2</sub> | 3 | 3.36 | 0.340 | 7.65 | 0.054 |
| block x species | 9 | 3.27 | 0.953 | 27.78 | **0.0010** |
| block x treatment | 3 | 6.62 | 0.085 | 3.09 | 0.378 |
| rosette surface at t<sub>2</sub> x species | 3 | 3.17 | 0.366 | 6.20 | 0.102 |
| rosette surface at t<sub>2</sub> x treatment | 3 | 0.69 | 0.407 | 16.81 | **4.13e-05** |
| species x treatment | 3 | 0.92 | 0.819 | 55.74 | **4.77e-12** |
| Accession (random) | 1 | 25.89 | **3.62e-07** | 377.7 | **< 2e-16** |

Then we analysed the four geographic areas within *C. bursa-pastoris*. We also detected a negative effect of competition on rosette surfaces but not on growth rate (Table 4). In addition we found a significant area x treatment effect for the second measure of rosette surface (Figure 2 and Table 4). However, the effect of competition was weak with *Ic* varying from 0.92 in Middle East to 0.99 in China, Central Asia and Europe being intermediate with 0.95 and 0.97, respectively. The contrast analysis on interactions indicated that Middle East is significantly different from Europe and China but all other comparisons are not (Table S4). As for comparison among species, competition had a strong and differential impact among geographic areas on flowers number (Table 5 and Figure 2). However, the pattern was almost reverse to rosette surface. China was the most sensitive to competition with *Ic* = 0.54 and Middle East the least, *Ic* = 0.81, Central Asia and Europe being intermediate with 0.71 and 0.69, respectively (Figure 2 and Table S5). Middle-East, Europe and China are significantly different from each others: Central Asia is significantly different from China but not from Europe and Middle-East. Rosette surface was positively and significantly correlated with flowers number, as expected (Table 5). Therefore, this relation cannot explain the reverse pattern for the effect of competition on rosette surface and flower size. It is worth noting that, although significant, the differences are very weak for rosette surface compared to flowers number.

**Figure 2:**
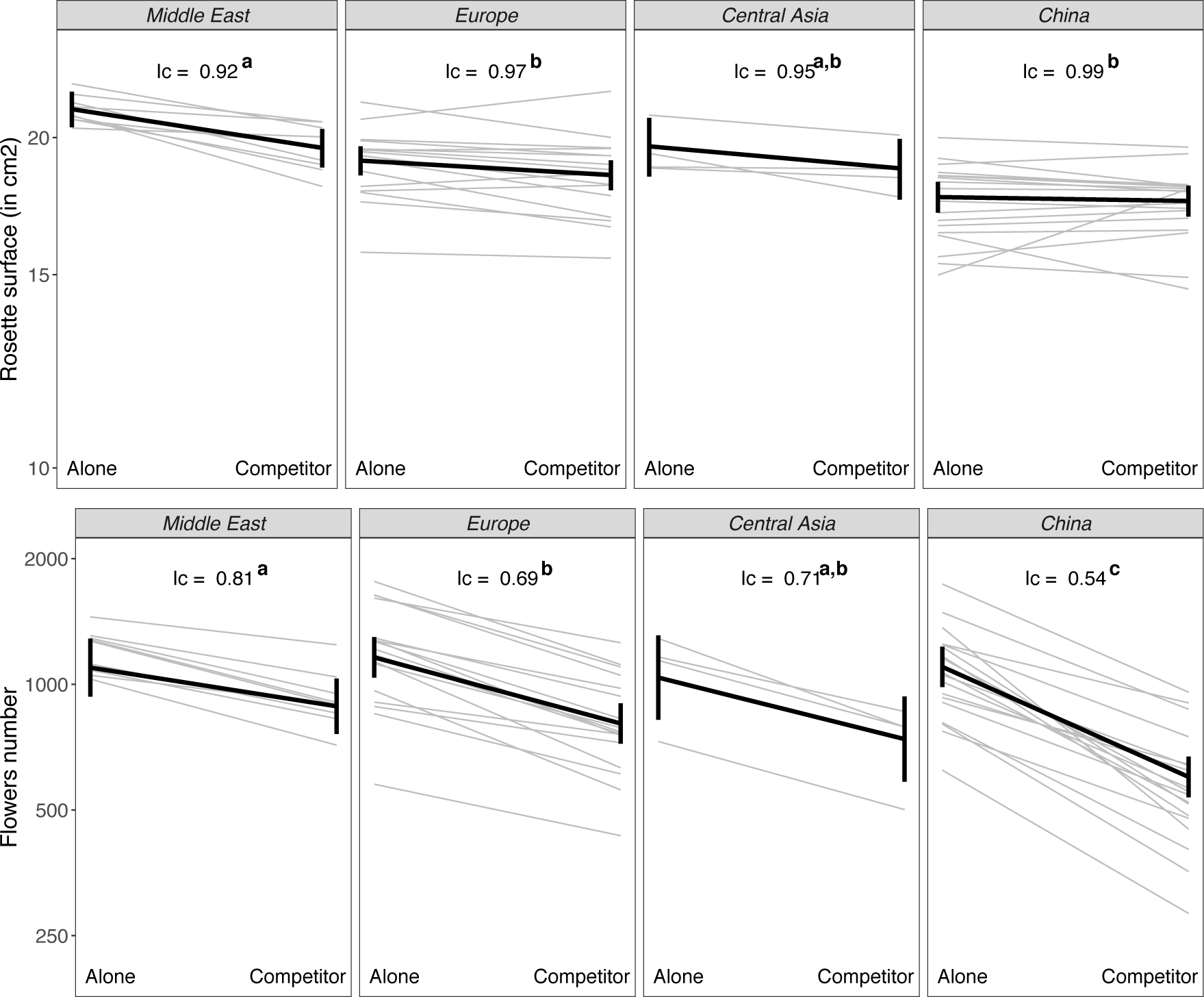
Rosette surface and flowers number without and with competitors for the four geographic area of *Capsella bursa + pastoris*. Same legend as in Figure 1.

**Table 4.** Analyses of variance for vegetative traits for the comparison among *C. bursa-pastoris* geographic areas. Same legend as Table 2

|  |  | Rosette surface 1 |  |  | Rosette surface 2 |  |  | Growth rate |  |  |
| --- | --- | --- | --- | --- | --- | --- | --- | --- | --- | --- |
|  | DoF | SS | F | p-value | SS | F | p-value | SS | F | p-value |
| block | 3 | 179.7 | 64.19 | **< 2.2e-16** | 94.41 | 27.51 | **8.88e-16** | 0.76 | 13.12 | **4.50e-08** |
| area | 3 | 19.01 | 6.79 | **0.0007** | 43.75 | 14.58 | **2.64e-06** | 0.08 | 1.40 | 0.254 |
| treatment | 1 | 17.67 | 18.94 | **1.85e-05** | 40.84 | 36.09 | **5.35e-09** | 0.02 | 1.22 | 0.270 |
| block x area | 9 | 5.62 | 0.70 | 0.708 | 8.76 | 0.89 | 0.536 | 0.24 | 1.49 | 0.152 |
| block x treatment | 3 | 1.12 | 0.41 | 0.740 | 2.01 | 0.61 | 0.607 | 0.15 | 2.73 | **0.044** |
| area x treatment | 3 | 6.57 | 2.46 | 0.063 | 26.20 | 7.96 | **3.95e-05** | 0.06 | 1.11 | 0.346 |
|  |  | LR |  | p-value | LR |  | p-value | LR |  | p-value |
| accession (random) | 1 | 177 |  | **< 2e-16** | 146 |  | **< 2e-16** | 70.4 |  | **< 2e-16** |

**Table 5.**
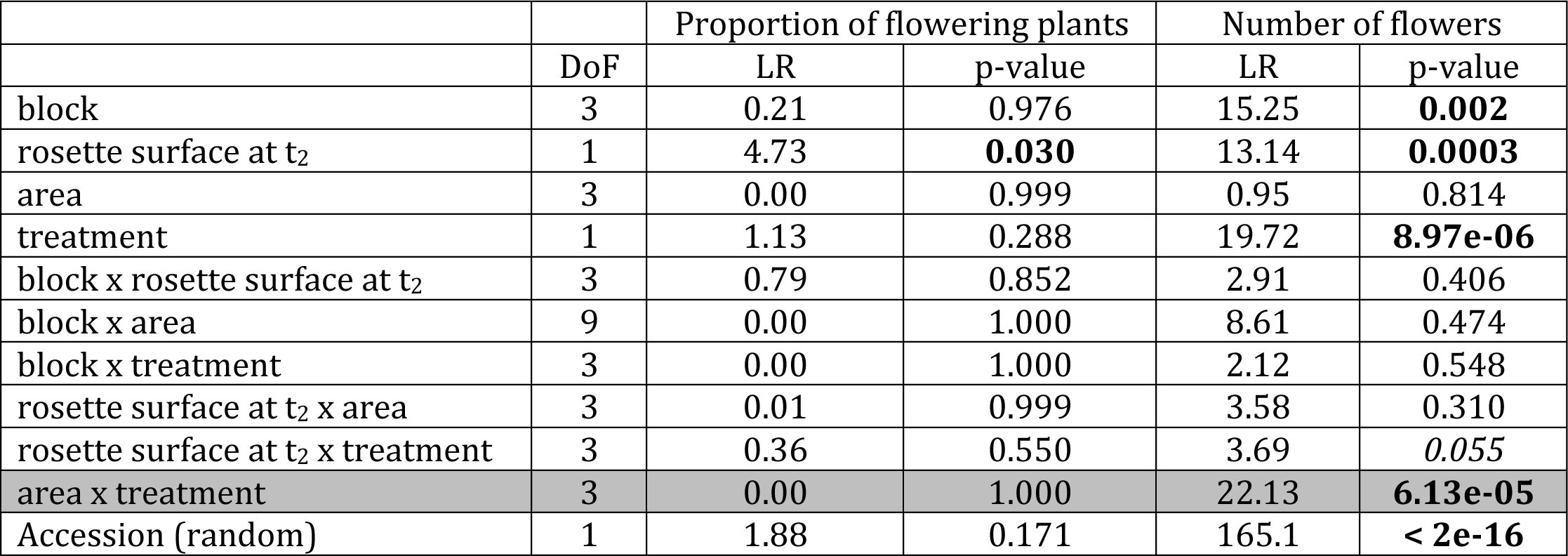
Analyses of deviance for reproductive traits for the comparison among *C. bursa-pastoris* geographic areas. Same legend as Table 2.

|  | DoF | Proportion of flowering plants |  | Number of flowers |  |
| --- | --- | --- | --- | --- | --- |
|  |  | LR | p-value | LR | p-value |
| block | 3 | 0.21 | 0.976 | 15.25 | **0.002** |
| rosette surface at t<sub>2</sub> | 1 | 4.73 | **0.030** | 13.14 | **0.0003** |
| area | 3 | 0.00 | 0.999 | 0.95 | 0.814 |
| treatment | 1 | 1.13 | 0.288 | 19.72 | **8.97e-06** |
| block x rosette surface at t<sub>2</sub> | 3 | 0.79 | 0.852 | 2.91 | 0.406 |
| block x area | 9 | 0.00 | 1.000 | 8.61 | 0.474 |
| block x treatment | 3 | 0.00 | 1.000 | 2.12 | 0.548 |
| rosette surface at t<sub>2</sub> x area | 3 | 0.01 | 0.999 | 3.58 | 0.310 |
| rosette surface at t<sub>2</sub> x treatment | 3 | 0.36 | 0.550 | 3.69 | 0.055 |
| area x treatment | 3 | 0.00 | 1.000 | 22.13 | **6.13e-05** |
| Accession (random) | 1 | 1.88 | 0.171 | 165.1 | **< 2e-16** |

Average genetic diversity decreases from Middle East (0.0023), Europe (0.0018) and to China (0.0015), which also corresponds to the decreasing order of competition indices. We thus tested more formally whether differences in genetic diversity could explain the observed difference among geographic areas. Genetic diversity had a positive and significant effect on rosette surface when it is used instead of population and area effects and the effect is significantly more positive without than with competitors (Table S6_A and Figure S1_A). Genetic diversity also had a positive effect on flowers number but the effect is only marginally significant (Table S6_B and Figure S1_B) and there is no significant interaction with the treatment level. Globally, the effect of genetic diversity is much weaker than the effect of geographic area and is not sufficient to explain the effect of geographic area on the sensitivity to competition.

## Discussion

It has been recognized for a long time that the evolution of self-fertilization is often associated with a series of ecological traits, such as colonizing ability, annuality or weediness. These ideas were recently tested more formally with a meta-analysis, using Grime’s CSR theory of ecological strategies: selfing was found associated with ruderal habit and lower competiveness (Munoz *et al.* 2016). To understand better how competitive ability may vary with mating system we compared the effect of competition in four *Capsella* species differing by their mating system and ploidy level. In the tetraploid selfer, *C. bursa-pastoris*, we also characterized the effect of competition across the expansion range of the species. Rapid range expansion thanks to good colonizing abilities is also an ecological attribute of selfing species (Randle *et al.* 2009; Grossenbacher *et al.* 2015), and has been suggested to be negatively associated with competitiveness (e.g. Burton *et al.* 2010).

### Competitive ability varies within and among *Capsella* species

Using the number of flowers as a proxy for fitness, our results are in agreement with theoretical predictions: the outcrossing species, *C. grandiflora*, was the least sensitive to competition while the two diploid selfers, *C. orientalis* and *C. rubella*, were the most sensitive (Figure 2). The tetraploid selfer, *C. bursa-pastoris*, was intermediate between its two putative parents (Figure 2), and within the species, the sensitivity to competition increased from the possible region of origin to the expansion front (Figure 4). In a previous experiment, *C. rubella* was already found as the most sensitive to competition but no significant difference was found between *C. grandiflora* and *C. rubella* (Petrone Mendoza *et al.* 2018). However, in that experiment only Greek accessions were used for *C. bursa-pastoris*, mainly corresponding to the Middle East genetic cluster (Cornille *et al.* 2016). Here too, Middle East accessions of *C. bursa-pastoris* (*Ic* = 0.81, Figure 2) did not differ from *C. grandiflora* (*Ic* = 0.74, Figure 1). The global difference between the two species was mainly attributable to the rest of the species range.

During the course of the experiment we also measured the surface of the rosette of the plants at two different times. While this trait is affected by competition, the global effect was rather weak and did not vary among species. Within *C. bursa-pastoris*, however, the effect of competition decreased across the species range, contrary to expectations and to what was observed for flower production. However, for both treatments, the raw values of rosette surface corresponded to the hypothesized qualities of the four geographic areas, with decreasing values from Middle East to China (Figure 2). Similarly, rosette surface increased with genetic diversity for both treatments, but more strongly without competitors such that the effect of competition is slightly increased with genetic diversity (Table S5 and Figure S1). Difference in absolute measures of vegetative traits, but not the effect of competition, are thus in agreement with the range expansion hypothesis. A possible caveat that could explain the difference between vegetative and reproductive traits is that rosette surface estimates were less precise than flowers number estimates. In addition, rosette surface is a less integrated fitness component than flowers number. Alternatively, although rosette surface is positively correlated with flowers number (Table 5) this could reflect the fact that rosette surface is not differentially affected by competition in this experiment. Indeed, access to light was likely not the main driver of competition in the experiment as differences in rosette surface could not explain (or if it did, only very weakly) the difference in flowers number between treatments (Table 5). In the experiment, most of the competition likely occurred at the root levels as we purposefully chose rather small pots. Unfortunately, underground biomass is very difficult to measure when roots are highly entangled. A more detailed analysis of functional traits remains to be done to understand how competition is mediated and whether there is any difference between selfing and outcrossing species.

The underlying causes of the observed pattern of competition on flowers number cannot be established from this sole experiment and the results are in agreement with two main theoretical (and non exclusive) hypotheses: the colonization/competition trade-off and the mutation load hypotheses. If transition to selfing is associated with transition to a more ruderal strategy and colonization of lower competitive habitats, relaxed selection on competitive traits could explain the stronger effect of competition on the three selfing species. The same rationale would apply for the increased sensitivity to competition across the expansion range in *C. bursa-pastoris*. Alternatively, competitive ability could be affected by the mutation load. In agreement with theoretical predictions on the genetic effects of selfing, selection against deleterious mutations is relaxed in the three selfing species compared to the outcrossing one (Brandvain *et al.* 2013; Slotte *et al.* 2013; Douglas *et al.* 2015). Moreover, within *C. bursa-pastoris*, Chinese populations corresponding to the expansion front showed evidence of a higher load than European and Middle East populations, which could also be explained by introgression from *C. orientalis* (Kryvokhyzha *et al.* submitted). So both among and within species, the mutation load is negatively correlated with competitive ability. As in other studies (reviewed in te Beest *et al.* 2012), we observed a difference between the diploid and the tetraploid selfers, the latter having a higher competitive ability than the former. This is also in agreement with the mutation load theory with a masking effect of deleterious mutations in polyploids (Comai 2005). Alternatively, it could be a direct consequence of increased cell size induced by genome doubling (Comai 2005; te Beest *et al.* 2012) or simply that *C. bursa-pastoris* has intermediate characteristics between is two parental species. The link with the colonization/competition trade-off hypothesis is less clear but we can speculate that gene redundancy could more easily reduce the pleiotropy of genes affecting both colonization and competition traits.

### Limits of the study and further directions

Our main results for flowers number are in agreement with theoretical predictions on the effects of mating system (including range expansion dynamics) and ploidy level on competitive abilities. They are also congruent with a meta-analysis showing association between outcrossing and competitiveness (Munoz *et al.* 2016).

However, there are several limitations to our study. First, we are not aware of other direct comparison among closely related species and our results could be only specific to the *Capsella* genus. In addition, we only compared four species, which is insufficient to draw general conclusions. Such results remain to be confirmed at a larger scale in different species. A possible design would be to replicate the experiment with several tens of pairs of sister (or closely related) species with contrasted mating systems, but other life-history traits being as similar as possible. Alternatively, the effect of competition could be estimated at a whole clade scale with frequent transitions in mating system and with an appropriate phylogenetic control. Similarly, such designs could be used to test more extensively the effect of polyploidization. Such experiments seem promising but demanding, as they would require proper fitness measures on several thousands or tens of thousand of plants.

Another limit is that we found no effect or slight effects in the opposite direction for vegetative traits. In a previous study (Petrone Mendoza *et al.* 2018), no difference in competitive ability among species was observed for vegetative traits. A better understanding of the eco-physiology of the species and more tuned experiment on the competition conditions (e.g. number and nature of competitors, soils characteristics) would be useful. Experiments in the field would also give a more realistic answer about the possible effects of competition in the wild.

Finally, our results can be interpreted in the mutation load framework, which is well developed from a theoretical point of view (Agrawal 2010; Agrawal & Whitlock 2012; Peischl *et al.* 2013; Peischl & Excoffier 2015; Gilbert *et al.* 2017). However, we still lack direct evidence of relationships between variation in load and variation in fitness under various competitive conditions. Such experiments have been conducted in Drosophila to test for the effect of competition on the strength and softness of selection against specific deleterious mutations (Laffafian *et al.* 2010; Ho & Agrawal 2012). In *Capsella* species, the effect of the load at the whole genome scale can be estimated through genomic approaches (Kryvokhyzha *et al.* submitted)so that the effect of the genome-wide mutation load on competitive ability could be more directly evaluated.

### Implication for the long:term evolution of selfing species

Whatever the underlying causes, the relationship between mating system and competitiveness suggests new implications for the long-term evolution of selfing species. In particular, the existence of a relationship between mating system and competitiveness could help resolve the paradox of selfing species that appear ecologically and demographically successful in the short term (Grossenbacher *et al.* 2015), but an evolutionary dead-end in the longer term (Stebbins 1957; Igic & Busch 2013; Wright *et al.* 2013). Some ecological conditions, such as disturbed, temporary or newly opened habitats, can both favour the evolution of selfing through selection for reproductive assurance, and correspond to weakly competitive environments that would allow their persistence despite poor competitive ability. Interestingly, selection is expected to be less severe when interspecific competition is low. This is expected to allow mutations to build up with limited demographic consequences (Agrawal 2010; Agrawal & Whitlock 2012). However, selfing species would be trapped into these weakly competitive habitats. Increase in competition, for example during succession, would increase extinction risk by competitive exclusion, especially because the accumulated genetic load would have stronger demographic impact than under the initially less competitive conditions (Agrawal 2010; Agrawal & Whitlock 2012).

A recent meta-analysis suggested that selfing species could experience diminished niche breadth over time despite geographic expansion (Park *et al.* 2017). The authors suggested that it could be due to the lack of long-term adaptive potential and accumulation of deleterious mutations. Reduced competitive ability could also prevent their establishment in new habitats. The scenario proposed by Park at al. (Park *et al.* 2017) is related to ours where the decrease in available habitats over time would finally reduce geographic range after initial range expansion. This would provide an ecological scenario for the higher extinction rate in selfers underlying the dead-end hypothesis and the maintenance of outcrossing species through species selection (Goldberg *et al.* 2010; Igic & Busch 2013). More generally, it could also contribute to explain how selfing and asexual lineages could “senesce” in diversification rates, as recently proposed (Ho & Agrawal 2017). If species derived from outcrossing ancestors are rapidly trapped into restricted non-competitive habitats, subsequent newly formed selfing species (from already-selfing ancestors) would also inherit restricted ecological niches without benefiting from higher reproductive assurance compared to ancestors.

Finally, under the global scenario proposed above, polyploidy could buffer the negative effect of selfing and delay the extinction risk. The association between selfing and polyploidy (Barringer 2007; Robertson *et al.* 2011) could be due both to the facilitation of the shift to polyploidy by selfing (Rodriguez 1996; Rausch & Morgan 2005) and to the reduction of extinction risk in selfers by polyploidy.

Combining genetic, ecological and demographical approaches has already been advocated to understand the transition from outcrossing to selfing (Cheptou 2007; Cheptou & Schoen 2007). We suggest that this should also be a promising approach to a better understanding of the long-term fate of selfing species.

## Acknowledgements

For this project, SG was supported jointly by the French CNRS and the Marie Curie IEF Grant “SELFADAPT” 623486. The authors declare no conflict of interest.

